# A glial cell derived pathway directs regenerating optic nerve axons toward the optic chiasm

**DOI:** 10.1101/2024.10.15.618346

**Authors:** Beth M. Harvey, Melissa Baxter, Alexis M. Garcia, Michael Granato

## Abstract

After optic nerve injury, several retinal ganglion cell (RGC) intrinsic signaling pathways have been shown to enhance RGC survival and RGC axonal growth. In contrast, few extrinsic cues have been identified that guide regenerating RGC axons toward and across the optic chiasm. Here, we use live-cell imaging in larval zebrafish to show that regrowing RGC axons initiate growth toward the midline and extend along a trajectory similar to their original projection. From a candidate genetic screen, we identify the glycosyltransferase Lh3 (also referred to as Plod3) to be required to direct regrowing RGC axons toward the midline during active regeneration. Moreover, we show that transgenic *lh3* expression in *sox10+* presumptive *olig2*+ oligodendrocytes located near the optic chiasm restores directed axonal growth in *lh3* mutants. Finally, we find that mutants in *collagen 18a1 (col18a1),* a putative Lh3 substrate, display RGC axonal misguidance phenotypes similar to *lh3* mutants, suggesting that *lh3* may act through *col18a1* during regeneration. Combined, these data identify *lh3* as part of a glial derived molecular pathway critical for guiding *in vivo* regenerating RGC axons toward and across the optic chiasm.

## Introduction

As a component of the central nervous system, retinal ganglion cell (RGC) axons within the mammalian optic nerve display a poor regenerative capacity. Multiple factors contribute to the minimal regeneration in mammals, including massive injury-induced RGC death and overall limited axonal regrowth (Villegas-Pérez *et al*., 1993; Berkelaar *et al*., 1994; Goldberg *et al*., 2002). Several studies have identified a number of RGC intrinsic signaling pathways that enhance RGC survival and increase long range axonal growth after injury. For example, deletion of the transcriptional repressor KLF4 in RGCs increases axonal regrowth (Moore *et al*., 2009). Similarly, activation of inflammatory responses within the retina through injection of Zymosan and cAMP enhances cell survival and axonal regrowth (Leon *et al*., 2000; Goldberg *et al*., 2002; Yin *et al*., 2009). Moreover, deleting intrinsic factors in RGCs such as PTEN and TSC1, which function as negative regulators of the mTOR pathway, improves both RGC cell survival and axonal growth (Park *et al*., 2008; Smith *et al*., 2009; Norsworthy *et al*., 2017). However, enhancing RGC axonal growth by manipulating RGC intrinsic programs frequently results in axonal misguidance as axons project inappropriately from the optic tract before and at the optic chiasm during the initial stages of regeneration (Kurimoto *et al*., 2010; Luo *et al*., 2013; Vincent Pernet *et al*., 2013; V. Pernet *et al*., 2013; Berry, Ahmed and Logan, 2019). This demonstrates a need to identify molecular and cellular mechanisms that guide RGC axons as they regrow toward and across the optic chiasm during regeneration.

Following a Central Nervous System (CNS) injury in mammals, many non-neuronal cells create an extracellular environment that is largely inhibitory to axonal regrowth. The cellular and molecular barriers that form within a lesion site have long been attributed to an influx of inflammatory cells and the upregulation of inhibitory molecules not conducive for robust regeneration (Yiu and He, 2006; Fitch and Silver, 2008). Additional studies in mice have since revealed roles for microglia and oligodendrocytes in functional recovery and remyelination from CNS injury (Wang *et al*., 2020; Mendonça *et al*., 2021; Brennan *et al*., 2022). Yet due to the dominance of factors inhibiting axonal regrowth, the cellular and molecular mechanisms that guide regrowing RGC axons following axonal injury are largely unknown. Recently, spontaneous CNS regeneration has been described in two mammalian species, the naked mole rat and the spiny mouse, demonstrating that pro-regenerative environments that support robust CNS regeneration are not unique to the CNS of non-mammalian vertebrates (Park *et al*., 2017; Nogueira-Rodrigues *et al*., 2022). Therefore, model organisms that exhibit spontaneous robust CNS axonal regeneration, without an overwhelmingly inhibitory extracellular environment, provide unique opportunities to identify mechanisms that promote CNS regeneration.

We recently developed a rapid and robust optic nerve transection assay in larval zebrafish to take full advantage of its regenerative capabilities and optical transparency to study functional optic nerve regeneration *in vivo* (Harvey, Baxter and Granato, 2019, 2023). At just 5 days post fertilization (dpf), larval zebrafish already possess a functional visual system driving several visually guided behaviors, such as prey capture (Brockerhoff *et al*., 1995; Easter and Nicola, 1996; Gahtan, Tanger and Baier, 2005). Following complete transections of the RGC axons in 5 dpf larvae, we observe axonal degeneration, subsequent initiation of regrowth and reinnervation of the optic tecta by 72 hours post transection (hpt) (Harvey, Baxter and Granato, 2019). We have also shown that RGC axonal regeneration in larval zebrafish occurs independent of RGC death or neurogenesis, and that visual function is restored by 8 days post transection (Harvey, Baxter and Granato, 2019). Using this optic nerve transection assay, we performed a candidate genetic screen and identify Lh3, a collagen-modifying glycosyltransferase whose gene name is referred to as *lh3* or also *plod3*, to be critical for directing axonal regrowth toward the optic chiasm during optic nerve regeneration. We show that following optic nerve transection, *lh3* expression is upregulated near the site of axonal injury and that Lh3 functions during the time when active regeneration occurs, specifically during the early stages of regeneration as RGC axons are growing toward the midline. We also show that Lh3 expression in oligodendrocytes is required to direct early regrowing RGC axons across the midline along their original path. Furthermore, we show that mutants in *collagen 18a1 (col18a1),* a putative Lh3 substrate, display RGC axonal misguidance phenotypes similar to *lh3* mutants, indicating at a glial-derived molecular pathway in which Lh3 acts through Collagen18a1 to direct RGC axons. Combined, our results demonstrate a critical pro-regenerative *in vivo* role for oligodendrocytes during early optic nerve regeneration to direct axon regrowth toward and across the midline.

## Results

### Lh3 is required for growth across midline during regeneration

To identify molecular mechanisms that guide regrowing RGC axons across the optic chiasm during optic nerve regeneration, we performed a candidate genetic screen consisting of 35 genes that are expressed in the visual system or have known functions in axon growth and guidance. Specifically, we used a sharpened tungsten needle to transect the optic nerves of genetic mutants and their siblings at 5 dpf and assessed at 72 hpt whether RGC axons had regrown across the optic chiasm to reinnervate the optic tectum. One of the top hits from this screen was the gene *lh3*, which encodes a multidomain glycosyltransferase that post-translationally modifies collagen proteins (Heikkinen *et al*., 2000; C. Wang *et al*., 2002). RGC axon projections in *lh3* mutants cross the midline and project to the optic tecta in a manner indistinguishable to their siblings (Zeller *et al*., 2002). However, we took advantage of a previously validated transgene *Tg(hsp70:lh3-myc)*, which expresses a Myc-tagged Lh3 under the control of the *hsp70* heat-shock-activated promoter (Isaacman-Beck *et al*., 2015) to rescue various morphological deficits that would affect the survivability of the larvae for the duration of the regeneration experiments (Zeller and Granato, 1999; Taler *et al*., 2019) (Fig. 1A). At 5 dpf, uninjured RGC axons in both *lh3* siblings and mutants exit the eye and project across the optic chiasm to terminate in the optic tecta (Fig. 1B-E). Following optic nerve transection, axons in *lh3* siblings grow across the midline and reinnervate the optic tecta (Fig. 1F,H,J,L). In contrast, transected optic nerve axons in *lh3* mutants fail to cross the midline and instead project along aberrant trajectories, often along anterior and posterior paths or also dorsally or ventrally (Fig 1G,I,K,L, magenta arrowheads). These results reflect a functional role for Lh3 during optic nerve regeneration in directing RGC axons towards and across the midline at the optic chiasm.

**Figure 1.**
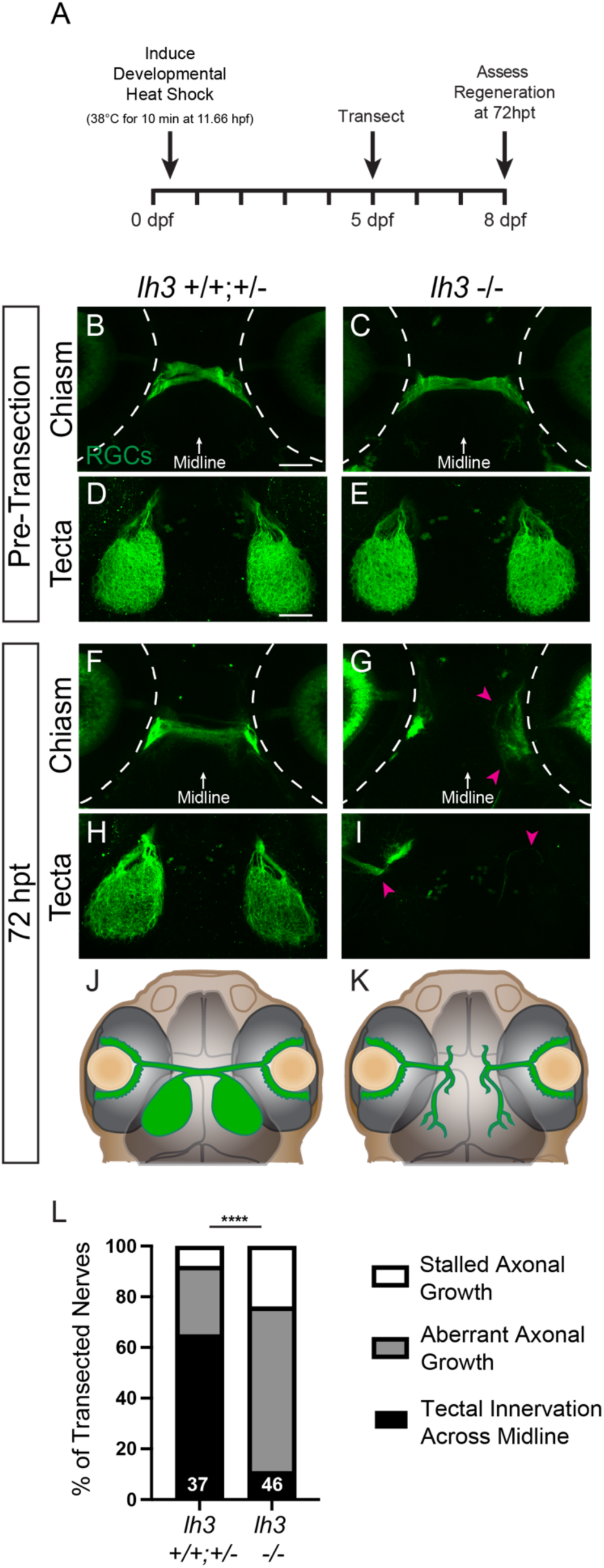
Lh3 is required for growth across the midline during regeneration. (A) Timeline of a transection experiment with *lh3* larvae (hpf, hours post fertilization; dpf, days post fertilization; hpt, hours post transection). (B-C) Before injury, RGC axons labeled by *Tg(isl2b:GFP)* in siblings and conditional *lh3* mutants cross the midline and (D-E) innervate contralateral tecta. (F-I) In contrast to *lh3* siblings at 72 hpt, RGC axons in *lh3* mutants turn away from the midline, project aberrantly (magenta arrowheads) and do not reinnervate contralateral optic tecta. (B-I) Dashed lines indicate the outline of the eyes; scale bars = 50 µm (J-K) Schematics of RGC axonal growth at 72 hpt for (J) *lh3* siblings and (K) *lh3* mutants. (L) Quantification of the percent of nerves at 72 hpt that grew across the midline to reinnervate the optic tectum, did not grow across the midline and instead grew along aberrant trajectories or showed stalled axonal growth. Numbers indicate total number of transected nerves. **** *P<0.0001*, Chi squared test. Representative images of chiasms and tecta for each timepoint are of the same fixed larva, though across timepoints are different larvae.

### *Lh3* expression is upregulated in the optic chiasm during regeneration

We next asked whether Lh3 is required during the process of optic nerve regeneration. For this, we took advantage of the conditional *Tg(hsp70:lh3-myc)* transgene (Isaacman-Beck *et al*., 2015). We induced Lh3 expression in 5 dpf larvae 4-5 hours prior to performing the optic nerve transections, and then assessed axonal regrowth across the optic chiasm and reinnervation of the optic tecta at 72 hpt (Fig. 2A). RGC axons in *lh3* siblings that did or did not receive a heat shock regrew across the midline and innervated the optic tecta (Fig. 2B,D,F). In contrast, *lh3* mutants without a heat shock display aberrant axonal growth, while heat shock induced Lh3 expression in *lh3* mutants restores optic nerve axonal regrowth across the midline (Fig. 2C,E-F). These data demonstrate that Lh3 acts during the process of optic nerve regeneration.

**Figure 2.**
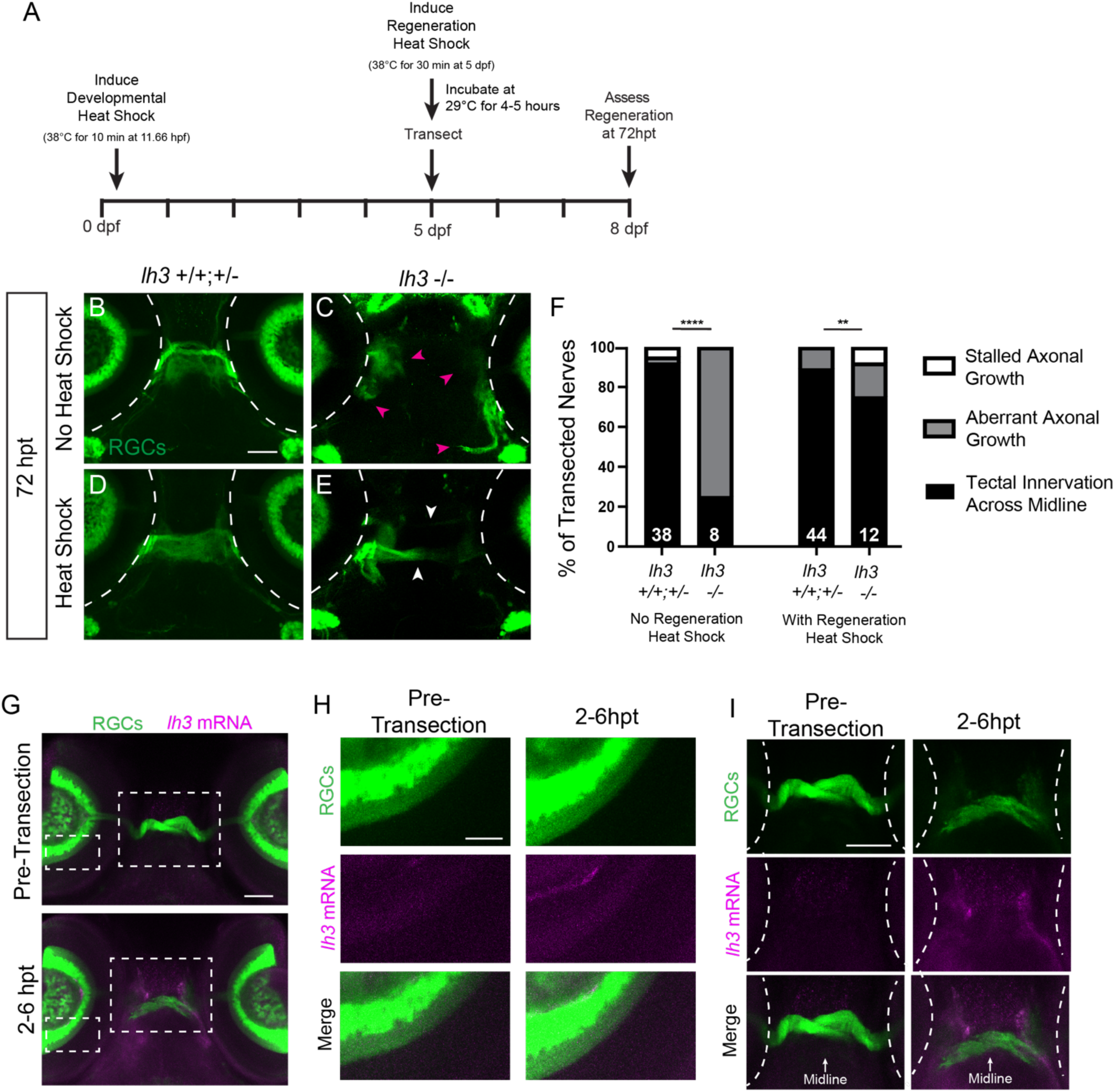
*Lh3* expression is upregulated in the optic chiasm during early regeneration. (A) Timeline of induced Lh3 expression with *Tg(hsp70:lh3-myc)* during development as well as regeneration (hpf, hours post fertilization; dpf, days post fertilization; hpt, hours post transection). (B-C) At 72 hpt in *lh3* siblings with no Heat Shock induction, RGC axons cross the midline to innervate the optic tecta, while RGC axons in *lh3* mutants turn away from the midline and project aberrantly (C, magenta arrowheads). (D-E) In *lh3* siblings and mutants that received a heat shock, RGC axons project across the midline (E, white arrowheads) and innervate the optic tecta. Dashed lines indicate the outline of the eyes; scale bar = 50 µm. (F) Quantification of the percent of nerves at 72 hpt that grew across the midline to reinnervate the optic tectum, did not grow across the midline and instead grew along aberrant trajectories or showed stalled axonal growth in *Tg(isl2b:GFP)* larvae with and without a regeneration heat-shock. Numbers indicate total number of transected nerves. \*\*\*\**P<0.0001*, \*\**P<0.01*, chi squared test. (G-I) *Lh3* mRNA expression in *Tg(isl2b:GFP)* larvae pre-transection and at 2-6 hpt. (G) White dashed boxes are areas of the retina shown in (H) and the optic chiasm shown in (I). Scale bars, (G) 50 µm, (H) 20 µm, (I) 50 µm. Axons in the chiasm region at 2-6 hpt have not yet fully degenerated following transection. Dashed lines indicate the outline of the eyes in (I). Representative images across timepoints are of different larvae.

We then sought to determine in which cell-type Lh3 was expressed during optic nerve regeneration. We used *in situ* hybridization chain reaction to detect *lh3* mRNA expression prior to and after optic nerve transections. Prior to transections, we failed to detect *lh3* mRNA expression in retinas nor in the region of the optic chiasm (Fig. 2G-I; n = 17 larvae). Similarly, after transection, we failed to detect *lh3* mRNA expression in RGC neurons in the retina (Fig. 2G-H; n = 41 larvae). In contrast, we detected upregulated *lh3* mRNA expression around the transected proximal stump of RGC axons and near the midline during early regeneration between 2 and 6 hpt (Fig. 2G,I; n = 30/41 larvae), demonstrating that *lh3* mRNA is upregulated following optic nerve transection in the region of the optic chiasm.

### Lh3 directs regrowing RGC axonal growth toward their original trajectory

Following optic nerve transections in wild-type larvae, we have previously observed new RGC axons emerging from the proximal stump of transected optic nerves by 24 hpt and exhibit robust growth across the optic chiasm by 48 hpt (Harvey, Baxter and Granato, 2019). To determine whether Lh3 continuously directs regenerating RGC axons once they emerge from the optic nerve stump, we used live cell imaging and examined the dynamics of regenerating RGC axons as they approach and cross the optic chiasm, which has not previously been reported. For simplicity, we removed one eye from 4 dpf larvae to only visualize the regrowth from a single retina per larva, and then performed transections to timelapse *lh3* siblings and mutants (Fig 3A, S1). Following transections on 5 dpf *lh3* sibling and mutant larvae, we observe that the portion of axons distal to the transection site undergoes Wallerian degeneration, as has been observed for other nerves in zebrafish (Fig. 3B-D,G-I, Supplementary Movies 1 and 2, brackets) (Waller, 1851; Martin *et al*., 2010; Rosenberg *et al*., 2012; Bremer *et al*., 2019). In *lh3* siblings, some axons emerge from the proximal stump of RGC axons in a range of directions, often extending and retracting rapidly (Fig. 3B-C, Supplementary Movie 1). By 16 hpt, we observe that a small group of axon fascicles grows toward the midline and rapidly extends across the midline (Fig. 3D-E, Supplementary Movie 1, white arrowheads). Over the next few hours, additional fascicles of axons are observed following the growth of the axons that have crossed the midline (Fig. 3F, Supplementary Movie 1) revealing *in vivo* axonal dynamics during the early stages of optic nerve regeneration. Axons in *lh3* mutants emerge at similar timepoints to *lh3* siblings from the proximal stump (Fig 3G-H, Supplementary Movie 2). However, unlike the *lh3* siblings, axonal growth does not persist across the midline and axons often grow anteriorly and posteriorly or dorsally and ventrally around the eye (Fig. 3I-K, Supplementary Movie 2, magenta arrowheads), strongly suggesting that during the early process of optic nerve regeneration, Lh3 directs the initial regrowth of RGC axons toward the midline.

**Figure 3.**
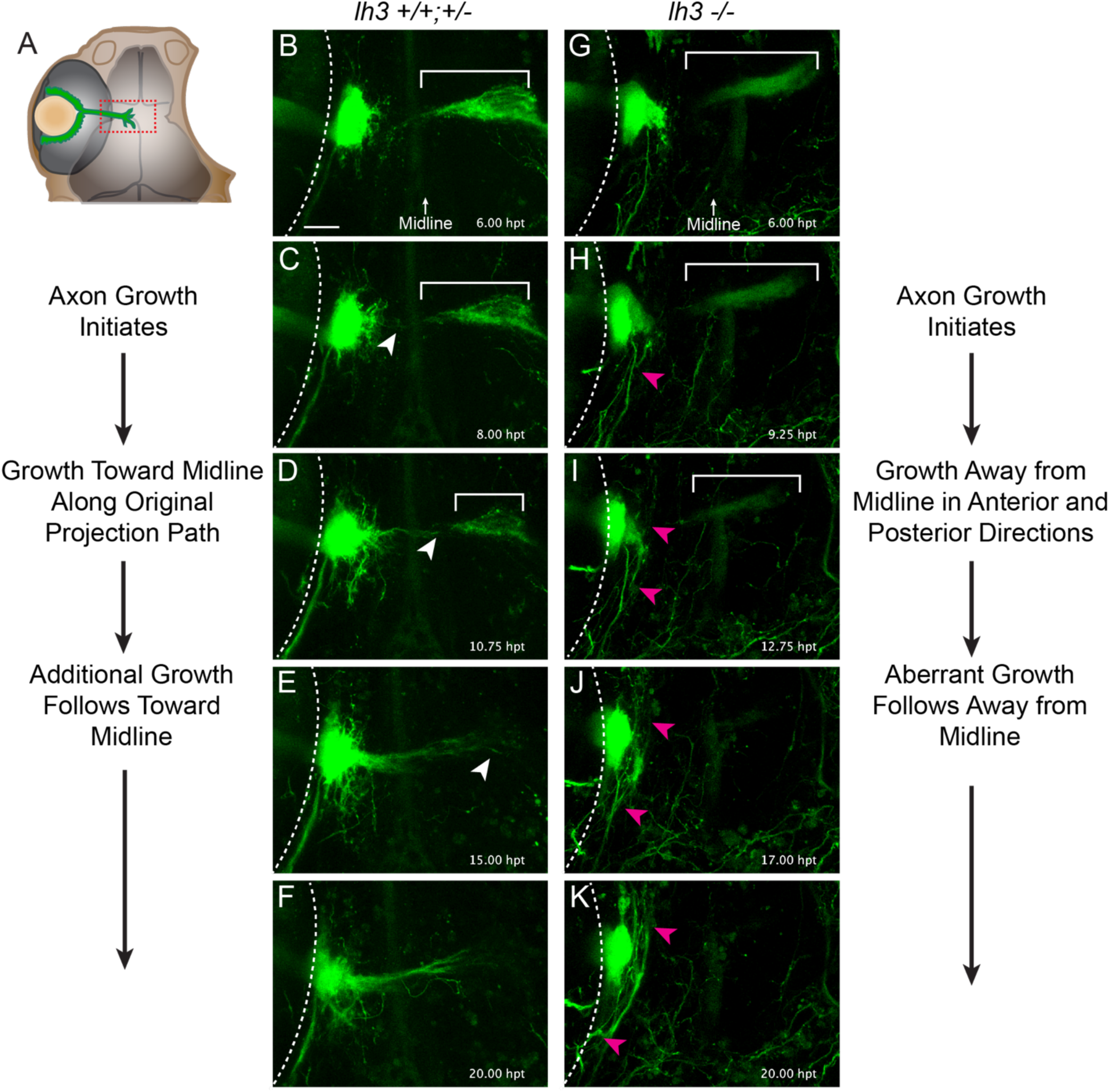
*Lh3* directs the initial regrowing RGC axons. (A) Schematic of larvae expressing *Tg(isl2b:GFP)* to label RGCs that were used for *in vivo* imaging. Larvae have one eye removed one day prior to timelapse imaging. Red box indicates the region imaged in panels B-K. (B,G) Transected RGC axons separate from the distal portion of the optic nerve that will eventually degenerate (indicated by brackets in B-D, G-I). (C) In *lh3* siblings, RGC axons initiate growth out from the proximal stump. (D) A small group of axons (white arrowhead) project directly toward the midline along the original path of the projection and then (E-F) subsequently additional axonal growth follows (n = 7/10 nerves). (H) In *lh3* mutants, axons grow out from the proximal stump, but (I-K) persist along aberrant trajectories (magenta arrowheads; n = 6/12 nerves). See also Figure S1. Dashed lines indicate the outline of the eyes; scale bar = 25 µm.

We next sought to identify the mechanisms by which Lh3 directs regrowing RGC axons toward and across the optic chiasm. We first asked whether Lh3 is critical for the initiation of axonal growth extending from the proximal stump of transected RGC axons, and consequently whether optic nerve regrowth was delayed in *lh3* mutants. We found no significant difference in the timing of initiation of axonal regrowth post transection (Fig 4A-B). To further define the trajectories that RGC axons take as they grow across the optic chiasm, we measured the angle between distinct axon fascicles and the original projection path before transection (Fig 4C; see Methods for more details). In optic nerves in *lh3* siblings and mutants that regrew across the midline, we observe that while some fascicles strayed from the pre-injury path, most fascicles grow very close to their original projection path (Fig 4D; Supplemental Table 1). Nerves in *lh3* siblings that did not grow toward the midline grew along a mostly single aberrant trajectory (Fig 4D; Supplemental Table 1). In contrast, in nerves that failed to cross the midline in *lh3* mutants, we found that the aberrant paths of axon projections displayed a much greater range of angles (Fig 4D; Supplemental Table 1). These data strongly suggest that *lh3* is required to direct RGC axons along their regenerative path, which is very similar to their pre-transection path, towards and across the optic chiasm.

**Figure 4.**
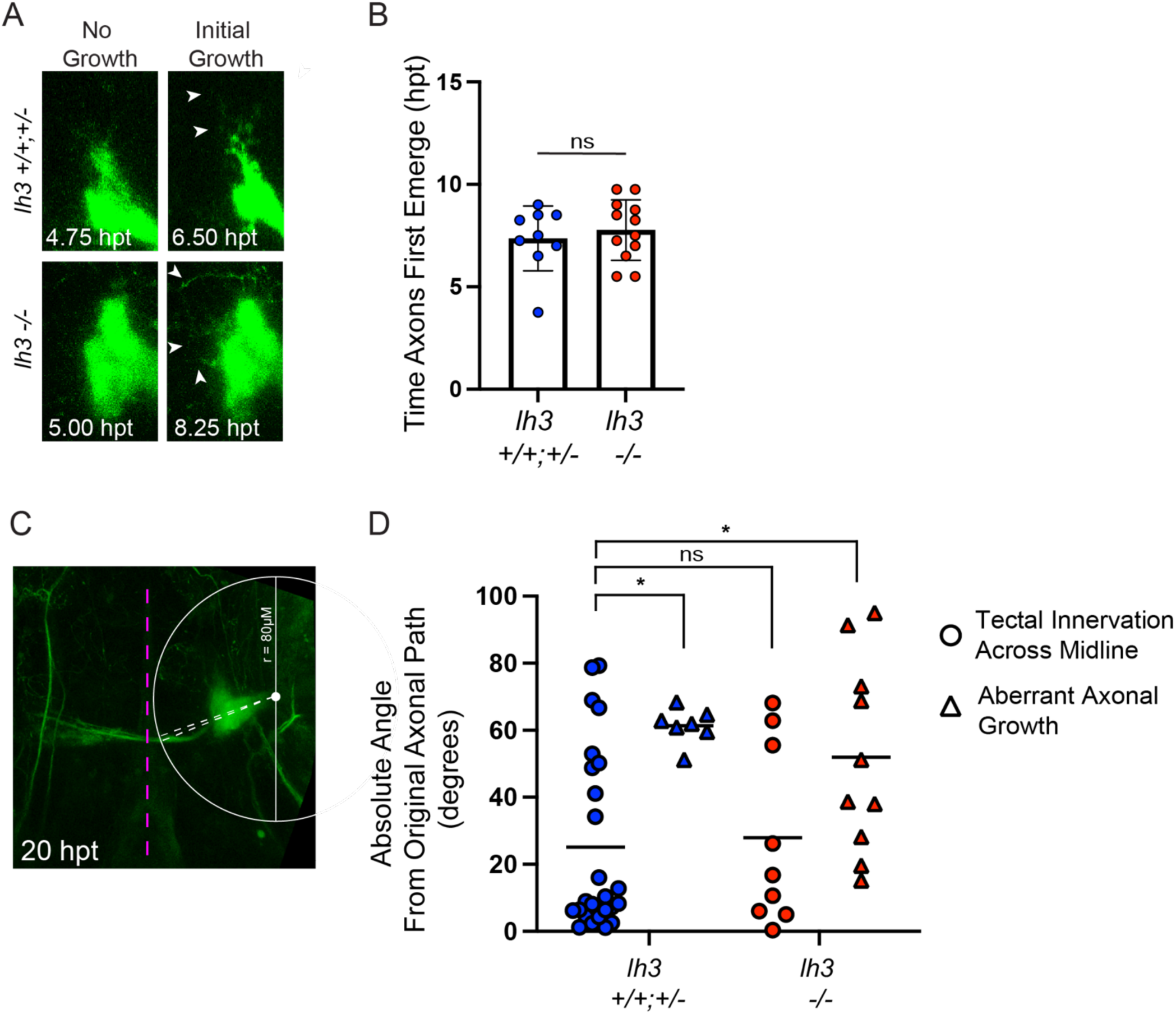
*Lh3* is required to direct regrowing RGC axons along their original path. (A) Representative frames from live timelapses of larvae expressing *Tg(isl2b:GFP)* to label RGCs showing the initial axonal growth that emerges from the injured proximal stump of axons. hpt = hours post transection. (B) Comparison of the time when axon fascicles first emerge from the proximal stump in *lh3* siblings and mutants. Time was determined by the first frame when axon fascicles were observed extending from the proximal stump that extended for at least one other consecutive frame. Data are represented as mean ± SD; n.s.*P>0.05*, t-test. (C) Representative image of a regenerating nerve at 20 hpt and the mark-up to measure the angle of growing fascicles from the injured stump of axons. The magenta line marks the parasphenoid cartilage. See Methods for more details on quantifications. (D) Quantification at 20 hpt of the absolute angle of axon fascicles from the end of a projection to the path of the original axonal projection. (Tectal Innervation Across Midline, sibling n = 6 nerves, mutants n = 3 nerves; Aberrant Axonal Growth, sibling n = 3 nerves, mutants n = 6 nerves). Individual dots are measured fascicle absolute angles, lines represent means; one-way ANOVA, n.s.*P>0.05*, * P<0.05. See Supplemental Table 1 for individual angles.

### Glial expression of Lh3 restores RGC axonal growth across the midline

Given that *lh3* mRNA expression is upregulated in area of the optic chiasm (Fig. 2I) and that *lh3* is required *in vivo* to direct regenerating RGC axons along their original projection path (Fig. 2F, 3, 4D), we hypothesized that the relevant cell type critical for *lh3* mediated optic nerve regeneration would be located in the region of the optic chiasm. Transcriptomics studies in mice have found *lh3* to be expressed most greatly in microglia (Zhang *et al*., 2014). Moreover, macrophages and microglia have been shown to clear debris from injured axons and can promote axonal growth and remyelination during CNS regeneration (Rosenberg *et al*., 2012; Wang *et al*., 2020; Brennan *et al*., 2022). Therefore, we first focused on macrophages and microglia as a potential source of Lh3 activity during optic nerve regeneration. In untransected larvae, we observe sporadic *mpeg+* cells near the optic chiasm (Fig 5A). At 24 hpt, we observe a greater concentration of *mpeg*+ cells surrounding the proximal stump of RGC axons and the new axonal growth (Fig. 5A; n = 11 larvae). We next used a transgenic rescue approach to test whether expression of *lh3* in macrophages/microglia is sufficient to restore axonal regrowth across the optic chiasm in *lh3* mutants. We found that Lh3 expression from a *Tg(mpeg:lh3-myc)* transgene fails to rescue the *lh3* mutant phenotype, where nerves in *lh3* mutants mostly display aberrant growth similar to *lh3* mutants without the *Tg(mpeg:lh3-myc)* transgene (Fig 5D). Since RGC axons regrow very closely to their original path, we next sought to examine oligodendrocytes associated with the optic nerve (Brösamle and Halpern, 2002). Using both a *Tg(olig2:dsred)* transgenic line labeling oligodendrocyte lineage cells (Fig 5B, n= 5 untransected larvae, n = 21 transected larvae) as well as *in situ* hybridization chain reactions for *mbpa* mRNA in myelinating oligodendrocytes (Fig 5C, n= 6 untransected larvae, n = 5 transected larvae), we observed oligodendrocytes in the optic chiasm and associated with RGC axons before and after transection (Kucenas *et al*., 2008; Ackerman and Monk, 2016). We then tested whether expression of *lh3* in oligodendrocytes would rescue axonal regrowth across the optic chiasm in *lh3* mutants. *Sox10*, like *olig2,* is a transcription factor that mediates oligodendrocyte lineage specificity and is maintained in all oligodendrocyte lineage cells (Bergles and Richardson, 2015; Ackerman and Monk, 2016). We find that *lh3* expression from a *Tg(sox10:lh3-mkate)* transgene restores RGC regrowth across the midline in otherwise mutant *lh3* larvae (Fig 5E). Furthermore, we detect *lh3* mRNA expression that overlaps with transgenic labeling of oligodendrocyte cells, demonstrating that *lh3* is upregulated in oligodendrocytes surrounding the proximal stump of transected RGC axons and along an axonal projection path across the midline during optic nerve regeneration (Fig. 5F-G, Supplemental Movie 3; n = 15/21). Altogether, these findings demonstrate that *lh3* expression in oligodendrocytes near the proximal stump of transected RGC axons directs early regrowing RGC axons toward the midline along their original path to reinnervate the optic tecta.

**Figure 5.**
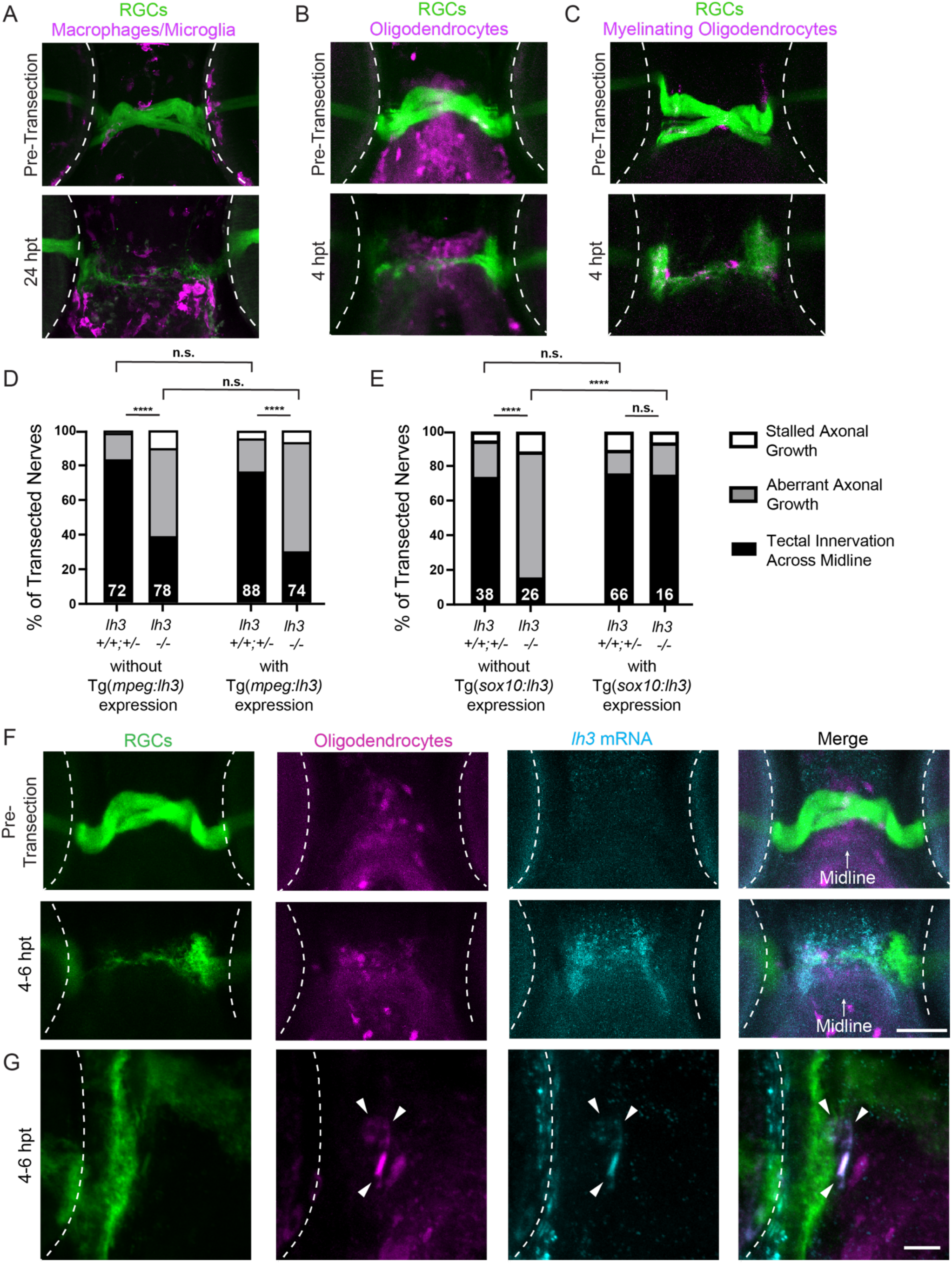
Lh3 in *sox10* expressing glia restores RGC axonal growth across the midline. (A) In wild-type larvae expressing *Tg(isl2b:GFP)* to label RGCs, macrophages and microglia labeled by *Tg*(*mpeg1:Gal4FF*);*Tg*(*UAS:nfsB-mCherry*) are observed in the area of the optic chiasm before transection. At 24 hours post transection (hpt), an influx of *mpeg*+ cells is seen. (B-C) Oligodendrocyte lineage cells labeled by *Tg(olig2:dsred)* as well as *mbpa* mRNA expression in myelinating oligodendrocytes are also seen in the chiasm before and after transection, near the injured proximal stump of RGC axons or along the degenerating original projection. (D) Quantification of the percent of nerves at 72 hpt that grew across the midline to reinnervate the optic tectum, did not grow across the midline and instead grew along aberrant trajectories or showed stalled axonal growth with and without *Tg(mpeg:lh3-myc)* transgene. Numbers indicate total number of transected nerves. \*\*\*\**P<0.0001*, n.s. *P>0.05*, chi squared test. (E) Quantification of regeneration with and without *Tg(sox10:lh3-mkate)* transgene. Numbers indicate total number of transected nerves. \*\*\*\**P<0.0001*, n.s. *P>0.05*, chi squared test. (F) *Lh3* mRNA expression at 4-6 hpt is upregulated and overlaps with oligodendrocytes labeled by *Tg(olig2:dsred)*. scale bars = 100um (G) Additional images of individual oligodendrocytes labeled by *Tg(olig2:dsred)* co-expressing *lh3* mRNA (see also Supplementary Movie 3). scale bars = 10um. Representative images across timepoints are of different larvae. Axons in the chiasm region at 4-6 hpt have not yet fully degenerated following transection. Dashed lines indicate the outline of the eyes.

### Col18a1 is also critical for directing regenerating RGC axons

We further sought to identify the relevant Lh3 substrate during optic nerve regeneration. Given the well documented role of Lh3 in post-translationally modifying collagen proteins (Heikkinen *et al*., 2000; C. Wang *et al*., 2002), we focused on collagen substrates. As part of our initial candidate genetic screen, we examined *col4a5*, which is required for proper RGC axon targeting of tectal layers and also functions to scaffold guidance cues during both the development of the visual system and during motor nerve regeneration (Xiao and Baier, 2007; Xiao *et al*., 2011; Isaacman-Beck *et al*., 2015; Murphy, Isaacman-Beck and Granato, 2022). We also tested *col7a1,* which has been shown to be expressed in the eye (Wullink *et al*., 2018). We ultimately found that RGC axon regeneration toward and across the midline is unaffected in *col4a5* and *col7a1* mutants (Fig. S2). We then examined *col18a1*, a collagen associated with Knobloch syndrome in humans that is characterized by eye abnormalities (Fukai *et al*., 2002; Suzuki *et al*., 2002). Col18a1 has several putative Lh3 sites within X-Lys-Gly-motifs and has been shown *in vitro* to bind to Lh3 (Risteli *et al*., 2004; Salo and Myllyharju, 2021). First, we used *in situ* hybridization chain reaction to detect *col18a1* mRNA expression and observe an upregulation of *col18a1* mRNA at 12 hpt in the optic chiasm, similar to the expression pattern of *lh3* following optic nerve transection (Fig. 6A; n = 13/14 larvae). We also detect *col18a1* mRNA expression that overlaps with transgenic labeling of oligodendrocyte cells, which demonstrates that *col18a1* is also upregulated in oligodendrocytes surrounding the proximal stump of transected RGC axons and along axonal projection path across the midline during optic nerve regeneration (Fig. 6B, Supplementary Movie 4). We then tested *col18a1* mutants in our optic nerve transection assay. Prior to transection, RGC axonal projections were indistinguishable between *col18a1* siblings and mutants (Fig. 6C-F). In contrast, at 72 hpt we observed regrowing RGC axons projecting away from the CNS midline, similar to what we observe in *lh3* mutants where regenerating RGC axons grow along aberrant paths (Fig 6G-K). This demonstrates that during regeneration *col18a1* is critical for directing regenerating RGC axons toward the midline and suggests that glial-derived Lh3 acts, as least in part, through Col18a1 during optic nerve regeneration.

**Figure 6.**
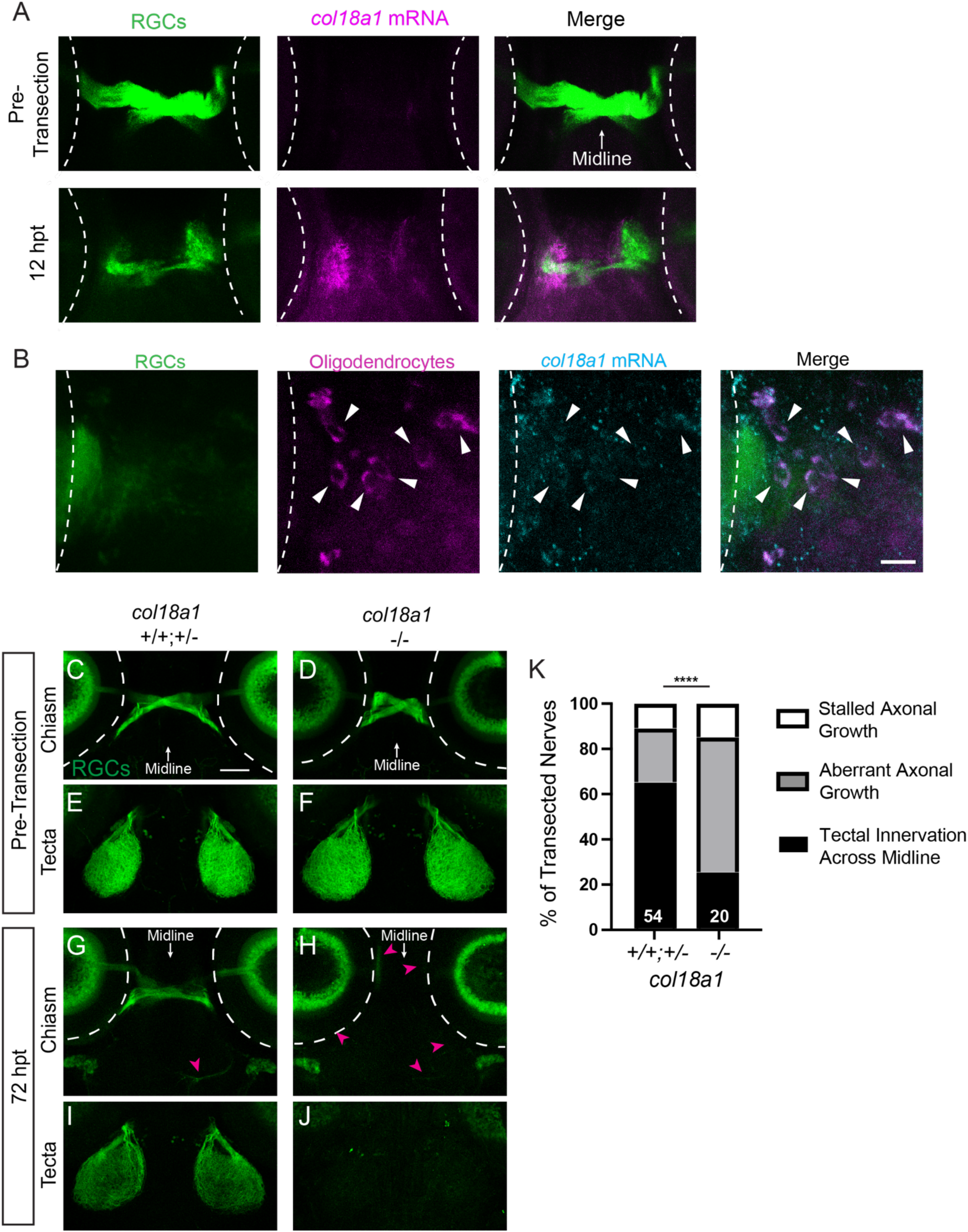
The Lh3 candidate substrate Col18a1 is upregulated after transection and directs RGC axons across the midline during regeneration. (A) *Col18a1* mRNA expression pre-transection and at 12 hours post transection (hpt) in larvae expressing *Tg(isl2b:GFP)* to label RGCs. Axons in the chiasm region at 12 hpt have not yet fully degenerated following transection. (B) Additional images of individual oligodendrocytes labeled by *Tg(olig2:dsred) co-expressing lh3* mRNA (see also Supplementary Movie 4). scale bars = 10um. (C-D) Before injury, RGC axons labeled by *Tg(isl2b:GFP)* in siblings and *col18a1* mutants cross the midline and (E-F) innervate contralateral tecta. (G-J) In contrast to *col18a1* siblings at 72 hpt, RGC axons in *col18a1* mutants turn away from the midline, project aberrantly (magenta arrowheads) and do not reinnervate contralateral optic tecta. Dashed lines indicate the outline of the eyes; scale bar = 50 µm. (K) Quantification of the percent of nerves at 72 hpt that grew across the midline to reinnervate the optic tectum, did not grow across the midline and instead grew along aberrant trajectories or showed stalled axonal growth. Numbers indicate total number of transected nerves. \*\*\*\**P<0.0001*, chi squared test. Representative images of chiasms and tecta for each timepoint are of the same fixed larva, though across timepoints are different larvae.

## Discussion

Regenerating axons often encounter multiple choice points along their paths that they must navigate properly to robustly reinnervate their original targets. Previous studies have shown that PTEN inactivation in RGC neurons strongly promotes RGC regrowth following an optic nerve injury, but that these axons display axonal misguidance at the optic chiasm, proving the optic chiasm to be one such choice point (Kurimoto *et al*., 2010; Luo *et al*., 2013; Vincent Pernet *et al*., 2013; V. Pernet *et al*., 2013; Berry, Ahmed and Logan, 2019). Here, we identify a mechanism critical for guiding RGC axons toward and across the optic chiasm during optic nerve regeneration. We find that *lh3* is required during regeneration to direct regenerating RGC axons toward and across the midline. We find that following optic nerve transection, expression of *lh3* and its presumptive substrate *col18a1* are upregulated in cells along the regenerative path near the optic chiasm. Using live cell imaging, we demonstrate that *lh3* is required to direct regrowing RGC axons along a regenerative trajectory very similar to their original path. Together, these results provide compelling evidence for a RGC extrinsic molecular pathway that guides optic axons during regeneration.

### Live cell imaging reveals axonal dynamics of early optic nerve regeneration

To date, most optic nerve regeneration studies in adult rodents and fish monitor RGC axonal regrowth with static time point analyses (Horder, 1971; Murray, 1976; V. Pernet *et al*., 2013; Yin and Benowitz, 2018). In contrast, peripheral nervous system regeneration, as well as spinal cord regeneration have been visualized in live larval zebrafish using *in vivo* imaging studies (Bhatt *et al*., 2004; Rosenberg *et al*., 2012; Ceci *et al*., 2014; Lewis and Kucenas, 2014; Isaacman-Beck *et al*., 2015; Bremer *et al*., 2019). To visualize RGC axonal dynamics during optic nerve regeneration, we previously established an optic nerve transection assay in larval zebrafish (Harvey, Baxter and Granato, 2019, 2023). Here, we perform live cell imaging of RGC axons initiating regrowth and regenerating across the optic chiasm *in vivo*. These timelapses reveal two intriguing insights into mechanisms that may mediate robust CNS regeneration in zebrafish. First, we observe that axons distal to the transection site undergo Wallerian degeneration (Fig. 3B-D,G-I, Movie S1 and Movie S2, brackets) (Waller, 1851). While a hallmark of the mammalian CNS is a slower rate of axonal fragmentation following axonal injury, here we observe complete degeneration of the distal axons by about 15 hpt (Vargas and Barres, 2007). While we did not examine whether persistence of degenerating debris would repel axonal growth, we do observe new axons reaching the midline around the same time axonal debris is cleared from that region (Fig. 3D). Axonal debris and associated myelin proteins are known to inhibit axonal regeneration (Vargas and Barres, 2007). But here, we find that Wallerian degeneration does not appear to be delayed in *lh3* mutants, and we therefore would not hypothesize axonal debris to contribute to the misguidance of RGC axons in *lh3* mutants.

Second, our live cell imaging reveals that a small group of axon fascicles exhibit strongly directed growth toward the midline by 16 hpt and subsequently extends across rapidly (Fig. 3D-E, Supplementary Movie 1, white arrowheads). Additional fascicles of axons continue to grow and follow the growth of the axons that initially crossed the midline (Fig. 3F, Supplementary Movie 1). These axons grow along a path that almost completely overlaps with their original path (Fig. 4D), and strongly suggests that there are factors expressed from cells associated with that projection that direct regrowing RGC axons during optic nerve regeneration.

### A novel role for oligodendrocytes in axon guidance during axon regeneration

To determine the relevant cell type through which *lh3* is functioning during optic nerve regeneration, we use cell-type specific transgenic rescue. Based on the *lh3* and *col18a1* mRNA expression in the optic chiasm (Fig. 2G-I, 6A), we hypothesized that the relevant cell type would be one found in that region during the initial stages of regeneration. We observe macrophages and microglia dynamically infiltrating the chiasm and surrounding regenerating axonal growth (Fig. 5A). *Tg(mpeg:lh3-myc)* expression fails to rescue aberrant axonal growth in *lh3* mutants, suggesting that *lh3* expression needs to be more specifically spatially regulated (Fig. 5D). We therefore examined oligodendrocytes and observe oligodendrocyte lineage cells present in the optic chiasm and we detect *mbpa* mRNA indicative of myelinating oligodendrocytes along the degenerating axons (Fig. 5B-C). We find that *lh3* expression from a *Tg(sox10:lh3-mkate)* transgene restores RGC regrowth across the midline in *lh3* mutants (Fig 5E). However, the transgenic tools we employed here do not allow is to distinguish whether oligodendrocyte precursor cells or mature oligodendrocytes were the relevant cell type for *lh3* and *col18a1,* as *sox10* and *olig2* are both transcription factors that are expressed in all oligodendrocyte lineage cells (Bergles and Richardson, 2015; Ackerman and Monk, 2016). Previous studies implicate oligodendrocytes to play roles in remyelination (Wang *et al*., 2020; Mendonça *et al*., 2021) and axon repulsion or inhibition via myelin-associated glycoprotein (McKerracher *et al*., 1994; Mukhopadhyay *et al*., 1994), NogoA (Chen *et al*., 2000; GrandPré *et al*., 2000), oligodendrocyte-myelin glycoprotein (K. C. Wang *et al*., 2002) and semaphorin 4D (Moreau-Fauvarque *et al*., 2003). From our results, we propose a mechanism in which Lh3 and Col18a1 expressed in oligodendrocytes mediate axon attraction to direct regenerating RGC axons along their original path, suggesting a novel role for oligodendrocytes in CNS regeneration.

### A molecular mechanism for *lh3* and *col18a1* in specifying a regenerative path

We identify *lh3* to be required to direct regenerating RGC axons toward and across the optic chiasm (Fig 1). We then determined that *col18a1* is also critical for directing RGC axons during optic nerve regeneration, but not another basement membrane *col4a5* or fibrillar *col7a1*, which demonstrates a specific role for *col18a1* in optic nerve regeneration (Fig 6, Fig S2). But the question remains: what are the mechanisms by which Lh3 and Col18a1 target RGC axons to their original path? We cannot exclude a possible mechanism involving RGC axons directly interacting with Col18a1 in the extracellular matrix through integrins (Elango *et al*., 2022). Transcriptomic analysis of RGCs at different stages of regeneration in adult zebrafish shows that several integrins are upregulated as axons grow toward and across the midline (Dhara *et al*., 2019). Therefore, the spatial and temporal specificity of Lh3 and Col18a1 expression could establish the regenerative path for regrowing RGC axons independent of any other guidance cues.

But in addition to playing structural roles, there is growing evidence showing a critical role for extracellular matrix proteins to bind and regulate the spatial distribution of axon guidance molecules in the extracellular space. For decades, glycosaminoglycans, such as heparan sulfate proteoglycans, have been known to regulate the extracellular localization of Slit repulsive guidance cues to affect axon guidance in development (Wang and Denburg, 1992; Johnson *et al*., 2004; Smart *et al*., 2011). More specifically, *col4a5* has been shown to be critical during development of the retinotectal projection for localizing Slit1 expression in the superficial layers of the optic tectum, which mediates lamina-specific axon targeting (Xiao and Baier, 2007; Xiao *et al*., 2011). During motor nerve regeneration, spatially restricted *col4a5* and *lh3* are required in a subset of Schwann cells to possibly create a Slit1a gradient that directs dorsal nerve axons to their original projection paths (Isaacman-Beck *et al*., 2015; Murphy, Isaacman-Beck and Granato, 2022). In this study, our results describe a similar mechanism where *lh3* is required to direct regrowing RGC axons toward the midline along their original path, but here, we find in optic nerve regeneration that Lh3 potentially acts through Col18a1, not Col4a5. What are possible guidance cues that Col18a1 could be spatially regulating? Col18a1 has three isoforms and within the long isoform is a Frizzled-like domain that has been shown to interact with Wnt molecules (Quélard *et al*., 2008). Moreover, Wnt signaling has been implicated in both promoting as well as repelling axonal growth in spinal cord and optic nerve regeneration (Yin *et al*., 2008; Hollis and Zou, 2012; Yam and Charron, 2013; Strand *et al*., 2016; Patel, Park and Hackam, 2017; Wehner *et al*., 2017). Therefore, we propose another potential mechanism in which Lh3 is required for Col18a1 expression that possibly spatially restricts Wnts in the extracellular space along the original RGC axonal projection to direct regenerating RGC axons during optic nerve regeneration. Altogether, our studies demonstrate that live cell imaging of optic nerve regeneration in larval zebrafish allows for unprecedented visualization of axonal growth dynamics and potential interactions with other cell types in the optic chiasm. Our data here reveal a glial derived molecular pathway that directs regenerating RGC axons across the optic chiasm during optic nerve regeneration, furthering our understanding of the cellular and molecular mechanisms that promote robust functional optic nerve regeneration.

## Methods

### Fish lines and maintenance

All zebrafish (*Danio rerio*) work was performed in compliance with the University of Pennsylvania Institutional Animal Care and Use Committee regulations and raised as previously described (Mullins *et al*., 1994). All transgenic lines were maintained in the Tupfel long fin genetic background except the *col18a1* line, which were maintain in the AB background. The following transgenic lines and mutants were used: *lh3^TV2O5^* (Schneider and Granato, 2006), *col18a1* (Isaacman-Beck *et al*., 2015), *col4a5^s510^* (Xiao and Baier, 2007), *col7a1^sa3473^*, *Tg(hsp:lh30-myc)* (Isaacman-Beck *et al*., 2015), *Tg(sox10:lh3-mkate)* (Isaacman-Beck *et al*., 2015), *Tg(isl2b:GFP)* (Pittman, Law and Chien, 2008), *Tg(olig2:dsred)* (Kucenas *et al*., 2008), *Tg*(*mpeg1:Gal4FF*)*^gl25^* (Ellett *et al*., 2011) and *Tg*(*UAS:nfsB-mCherry*)*^c264^* (Davison *et al*., 2007).

Adult zebrafish and embryos were raised at 28°C on a 14-h:10-h light:dark cycle, or 29°C for larvae in PTU E3, see Transection Assay.

To generate *Tg(mpeg1.1:lh3-myc)*, Gateway cloning was employed to combine *lh3-myc* into a pDest vector with the mpeg1.1 promoter and Tol2 transposon sites. Tol2 transgenesis was performed as previously described(Suster *et al*., 2009) into 1-cell stage embryos from *lh3* heterozygous incrosses. Transgenic lines were identified by genotyping for the transgene with forward primer ACAGTCTCTTGCGTCATCAAAACC and reverse primer TGATCACCAGCAGCTCATTG.

### Transection Assay

Optic nerve transection assays were performed as previously described(Harvey, Baxter and Granato, 2019, 2023). Briefly, to inhibit melanocyte pigmentation, larvae were raised in phenylthiourea (PTU, 0.2mM in E3 medium) in the dark at 29°C beginning at 1 dpf. For live cell imaging and distal transections, larvae were anesthetized in PTU E3 plus 0.0053% tricaine then mounted in 2.5% low-melt agarose (SeaPlaque, Lonza) prepared with PTU E3 plus 0.016% tricaine ventral-up on a glass microscopy slide. Then one eye was completely removed using forceps at 4 dpf. Larvae recovered in Ringer’s solution and then were kept in PTU E3 at 29°C until transected. At 5 dpf, larvae were anesthetized, mounted in agarose, and then optic nerve transections were performed on an Olympus SZX16 fluorescent microscope or a dissecting microscope with a NightSea Stereo Microscope Fluorescence Adapter (RB-GO only wavelength set, bandpass filter) using a sharpened tungsten needle (Fine Science Tools, Tip Diameter: 0.001mm, Rod Diameter: 0.125mm). Optic nerves were transected at the region of the nerve distal to it exiting the eye, yet proximal to the optic chiasm. Following injuries, larvae were removed from the agarose, allowed to recover in Ringer’s solution with 0.2mM PTU, then returned to 0.2mM PTU, E3 medium at 29°C. Larvae were inspected for transection efficiency at 16-18 hpt, except larvae that were fixed at early post transection timepoints, and only larvae with complete optic nerve transections with no visible intact axons remaining from the eye to the tectum were kept until fixation at later designated timepoints.

### Heat Shock-Induced *lh3* Rescue

To induce expression of Lh3 from *Tg(hsp:lh3-myc)* transgene, during development: Embryos were developmentally staged in the late afternoon. No more than 6 embryos were placed per tube in PCR strip-tubes. Tubes were incubated in a thermal cycler at 28°C until embryos would reach 11.66 hpf, then heated to 38°C for 10 minutes, then held at 28°C until they could be removed from the tubes.

To induce expression of Lh3 from Tg(hsp:lh3-myc) transgene, during regeneration: Larvae at 5 dpf homozygous for the Tg(hsp:lh3-myc) transgene were placed only one per tube in PCR strip-tubes. Tubes were incubated in a thermal cycler at 38°C for 30min then held at 29°C until they could be removed from the tubes. Larvae were transected 4-5 hours later.

### Immunostaining

Larvae were stained using methods modified from those previously described (Randlett *et al*., 2015). Briefly, larvae were fixed in 4% PFA in PBS overnight at 4°C. Larvae were washed in PBS + 0.25% Triton (PBT), incubated in 150mM Tris-HCl pH 9.0 for 15 min at 70°C, then washed in PBT. Larvae were permeabilized in 0.05% Trypsin-EDTA for 5 min on ice, washed in PBT, blocked in PBT containing 1% bovine serum albumin (BSA), 2% normal goat serum (NGS) and 1% dimethyl sulfoxide (DMSO), and then incubated in primary and secondary antibodies overnight at 4°C in PBT containing 1% BSA and 1% DMSO. Stained larvae were stored and mounted in Vectashield (Vector Laboratories) for imaging using confocal microscopy. For static timepoint analysis of *lh3* and *col18a1* mutants, primary antibodies used were mouse anti-GFP (JL-8, 1:200, Takara Bio), and secondary antibodies used were goat anti-mouse Alexa 488 (1:500, Molecular Probes). For static timepoint analysis of macrophages/microglia, primary antibodies used were mouse anti-GFP (JL-8, 1:200, Takara Bio), rabbit anti-RFP (1:200, Abcam ab62341) and secondary antibodies used were goat anti-mouse Alexa 488 (1:500, Molecular Probes) and goat anti-rabbit Alexa 594 (1:500, Molecular Probes). For static timepoint analysis of oligodendrocytes, primary antibodies used were rabbit anti-GFP (1:500, Molecular Probes), mouse anti-dsred (1:500, BD Pharmingen) and secondary antibodies used were goat anti-rabbit 488 (1:500, Molecular Probes) and goat anti-mouse Alexa 594 (1:500, Molecular Probes).

### Fluorescent *in situ* Hybridization with Hybridization Chain Reaction (HCR)

*Lh3 (plod3), col18a1 and mbpa* mRNA expression were detected using fluorescent *in situ* hybridization with HCR (Molecular Instruments, Los Angeles, CA, USA)(Choi *et al*., 2018). HCR probes, buffers, and hairpins were purchased from Molecular Instruments. *Tg(isl2b:GFP)* larvae were fixed at 5 dpf with 4% paraformaldehyde in PBS overnight at 4°C. No antibody staining was required together with in situ HCR. The staining was performed as previously described(Shainer *et al*., 2023).

### Static Confocal Imaging, Processing and Analysis

Larvae were imaged on a Zeiss 880 confocal microscope using a 20× objective. Airyscan super-resolution settings were used to obtain Figures 5G and 6B. Image stacks of optic chiasms or tecta were compressed into maximum intensity projections. All images were adjusted for brightness and contrast in Adobe Photoshop CS5, and color assigned using Fiji.

### Live Cell Imaging, Processing and Analysis

Larvae were anesthetized in PTU E3 with 0.0053% tricaine then mounted in 2.5% low-melt agarose (SeaPlaque, Lonza) prepared with PTU E3 plus 0.016% tricaine ventral-up on a glass microscopy slide. Then, a portion of the jaw, including the Meckel’s cartilage, palatoquadrate, and the most distal parts of the basihyal and ceratohyal structures, as well as one eye were carefully removed using forceps at 4 dpf. Care was taken so as not to damage the heart. Larvae recovered in Ringer’s solution and then were kept in PTU E3 at 29°C until transected. At 5 dpf, larvae were anesthetized again, mounted in agarose, and then optic nerve transections were performed. Larvae recovered in Ringer’s solution and then were kept in PTU E3 at 29°C until imaged for live cell imaging. For live imaging, larvae were mounted ventral side up in 35mm glass-bottomed Petri dishes in 1.5% low-melt agarose with 0.016% tricaine. Larvae were imaged using a Leica SP8 confocal microscope with the resonant scanner on, equipped with a petri dish heater and heated water perfusion so fresh E3 water flowed through the petri dish and the temperature was kept constant at around 28°C during the imaging session. Larvae were imaged from the ventral side of the brain, which was not covered by any jaw tissue. Image stacks were acquired every 10-15 minutes with a 20x 1.0 NA lens. All images were adjusted for brightness and contrast, and color assigned using Fiji.

To determine the time of growth initiation extending from the proximal stump of transected RGC axons, we recorded in which frame axonal extension longer than 1um occurred that continued to extend for another subsequent frame.

To measure the angle of the original path before transection to the end of distinct axon fascicles, we first rotated the images so that parasphenoid cartilage was precisely aligned vertically against a straight line drawn in Fiji. A circle with a radius of 80 uM centered where RGC axons exit the eye was drawn within Fiji using a Make Oval Macro in timelapse image sequences. Axons and fascicles terminating farther than the 80 uM region of interest at 20 hpt were used for measuring. First the angle of the original path to the parasphenoid was determined using frames before Wallerian degeneration was complete. Then the angle of the axons and fascicles to the parasphenoid were determined, and then the difference was calculated and graphed.

### Statistical Analysis

Data were imported into GraphPad Prism for analysis. Statistical analysis was performed as indicated throughout the text.

## Supporting information

Supplementary Movie related to Fig 3

Supplementary Movie related to Fig 3

## Acknowledgments

We would like to thank Dr. Andrea Stout of the UPenn Cell and Developmental Biology Microscopy Core, as well as Dr. Gordon Ruthel of the Penn Vet Imaging Core for providing technical assistance. We thank all members of the Granato laboratory for comments and discussion.

## Author Contributions

B.M.H. and M.G. conceptualized the experiments; B.M.H. M.B, and A.M.G. performed and analyzed experiments; B.M.H. wrote the original draft. All authors contributed to editing and revision of the paper.

## Data Availability

All relevant data are within the article and its supplementary information files.

## Supplemental Information

**Supplementary Movie 1**

Related to Figure 2, contains the movie which corresponds to the *lh3* sibling axonal regeneration time series shown in Figure 2B-F.

**Supplementary Movie 2**

Related to Figure 2, contains the movie which corresponds to the *lh3* sibling axonal regeneration time series shown in Figure 2G-K.

**Supplementary Movie 3**

Related to Figure 5G, representative three-dimensional rotation from a confocal z-stack demonstrating co-expression of *lh3* mRNA with *Tg(olig2:dsred)* oligodendrocytes.

**Supplementary Movie 4**

Related to Figure 6B, representative three-dimensional rotation from a confocal z-stack demonstrating co-expression of *col18a1* mRNA with *Tg(olig2:dsred)* oligodendrocytes.

**Supplemental Table 1**

Related to Figure 4, contains raw data of individual angles of fascicles from nerves that grew either across the midline or aberrantly from *lh3* sibling and mutant larvae.

---

**Figure S1.**
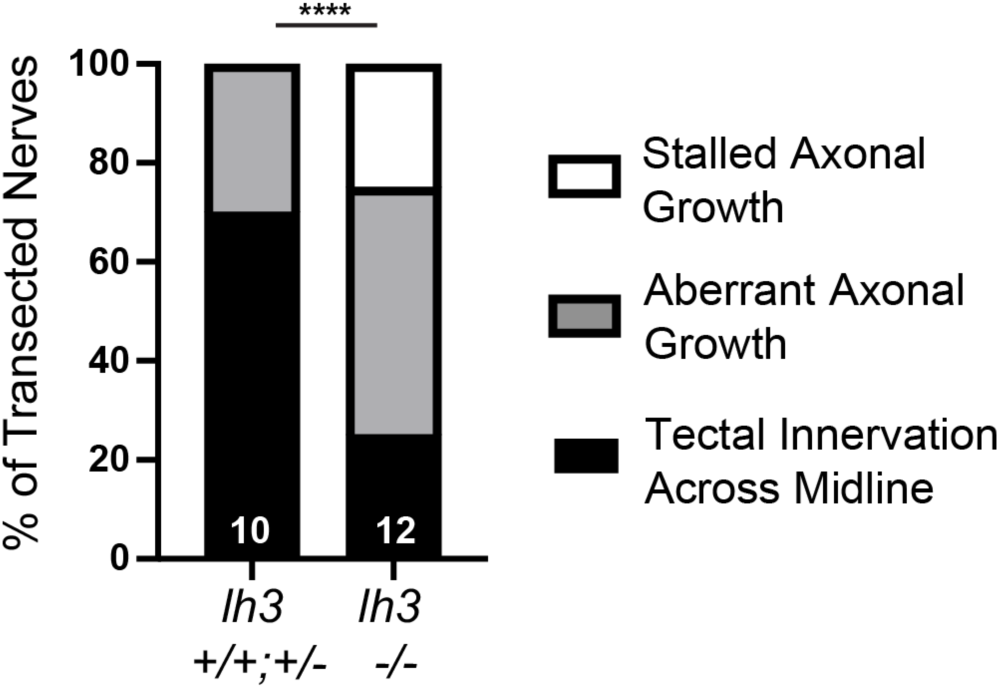
Related to Figures 3 and 4. Quantification of the percent of nerves from timelapsed larvae that grew across the midline to reinnervate the optic tectum, did not grow across the midline and instead grew along aberrant trajectories, or did not exhibit any axonal growth, ie. stalled, at 72 hpt. Numbers indicate total number of transected nerves. *P<0.00001*, Fisher’s exact test.

**Figure S2.**
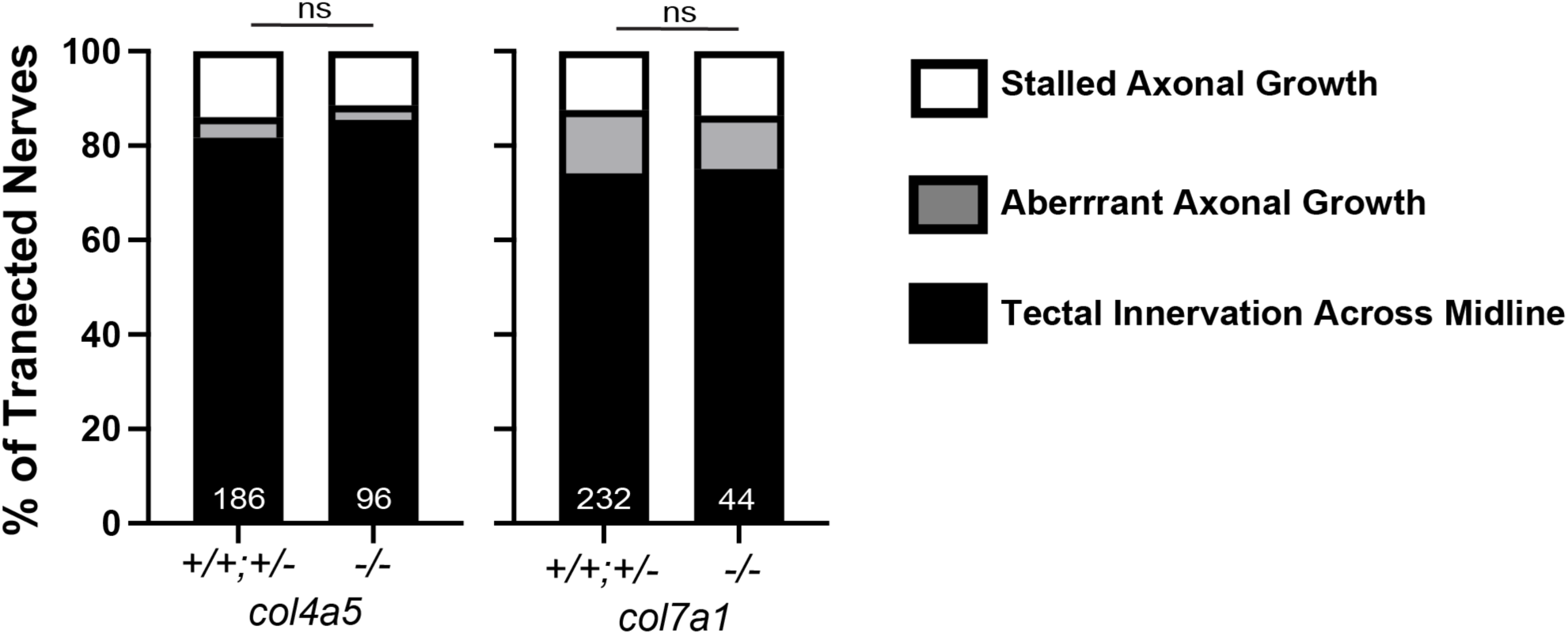
*Col4a5* and *col7a1* are not required to direct regrowing RGC axons to the midline during regeneration. Quantification of the percent of nerves that grew across the midline to reinnervate the optic tectum, did not grow across the midline and instead grew along aberrant trajectories, or did not exhibit any axonal growth, ie. stalled, at 72 hpt. Numbers indicate total number of transected nerves. n.s.*P>0.05*, chi-square test.

